# Genome-wide CRISPR Screens Reveal Genetic Mediators of Cereblon Modulator Toxicity in Primary Effusion Lymphoma

**DOI:** 10.1101/619312

**Authors:** Ajinkya Patil, Mark Manzano, Eva Gottwein

**Author notes:** **Corresponding Author:** Eva Gottwein, Department of Microbiology-Immunology, Northwestern University Feinberg School of Medicine, 320 E Superior St., Tarry Bldg., Room 6-735, Chicago, Illinois 60611, USA, e-mail address.

## Abstract

Genome-wide CRISPR/Cas9 screens represent a powerful approach to study mechanisms of drug action and resistance. Cereblon modulating agents (CMs) have recently emerged as candidates for therapeutic intervention in primary effusion lymphoma (PEL), a highly aggressive cancer caused by Kaposi’s sarcoma-associated herpesvirus. CMs bind to cereblon (CRBN), the substrate receptor of the cullin-RING type E3 ubiquitin ligase CRL4^CRBN^, and thereby trigger the acquisition and proteasomal degradation of neosubstrates. Downstream mechanisms of CM toxicity are incompletely understood, however. To identify novel CM effectors and mechanisms of CM resistance, we performed positive selection CRISPR screens using three CMs with increasing toxicity in PEL: lenalidomide (LEN), pomalidomide (POM), and CC-122. Results identified several novel modulators of the activity of CRL4^CRBN^. The number of genes whose inactivation confers resistance decreases with increasing CM efficacy. Only inactivation of CRBN conferred complete resistance to CC-122. Inactivation of the E2 ubiquitin conjugating enzyme UBE2G1 also conferred robust resistance against LEN and POM. Inactivation of additional genes, including the Nedd8-specific protease SENP8, conferred resistance to only LEN. SENP8 inactivation indirectly increased levels of unneddylated CUL4A/B, which limits CRL4^CRBN^ activity in a dominant negative manner. Accordingly, sensitivity of SENP8-inactivated cells to LEN is restored by overexpression of CRBN. In sum, our screens identify several novel players in CRL4^CRBN^ function and define pathways to CM resistance in PEL. These results provide rationale for increasing CM efficacy upon patient relapse from a less efficient CM. Identified genes could finally be developed as biomarkers to predict CM efficacy in PEL and other cancers.

**Key Points:**

1. Genome-wide CRISPR/Cas9 screens identify novel mediators of resistance to lenalidomide, pomalidomide and CC-122 in PEL cells.
2. UBE2G1 and SENP8 are modulators of CRL4^CRBN^ and their inactivation drives resistance to CMs in PEL-derived cell lines.

## Introduction

Primary effusion lymphoma (PEL) is a non-Hodgkin B cell lymphoma (NHL) caused by Kaposi’s Sarcoma-associated Herpesvirus (KSHV)^1,2^. PEL most commonly arises in HIV infected individuals, where it comprises ~4% of HIV-related NHLs^3^. PEL carries a poor prognosis, despite chemotherapy and anti-retrovirals^4,5^. Recent work by us and others suggests that cereblon modulators (CMs), including the immunomodulatory drugs (IMiDs) lenalidomide (LEN) and pomalidomide (POM), may present a promising treatment strategy in PEL^6,7^. LEN and POM are used in multiple myeloma (MM), as frontline therapy or upon relapse from LEN, respectively. Despite substantial efficacy of IMiDs in MM, patients commonly relapse. LEN is also effective in myelodysplastic syndrome with deletion of chromosome 5q [del(5q) MDS]. CC-122 is a fourth generation CM currently under pre-clinical investigation in diffuse large B cell lymphoma (DLBCL)^8^. The molecular target of CMs is cereblon (encoded by *CRBN*), a substrate receptor of the cullin-RING type E3 ubiquitin ligase CRL4^CRBN 9^.

The covalent conjugation of ubiquitin (Ub) to target proteins is mediated by an enzymatic cascade that involves an E1 Ub activating enzyme, E2 Ub conjugating enzyme, and E3 Ub ligase, which attaches Ub to lysine (K) residues of protein substrates or Ub itself^10^. K48-linked Ub chains of four or more moieties canonically target substrates for degradation by the 26S proteasome. The modular cullin-RING E3 Ub ligases (CRLs) are comprised of one of seven cullin family scaffold proteins (CUL1, 2, 3, 4A, 4B, 5, or 7), which recruits a RING family protein with E3 ubiquitin ligase function and a substrate recognition module^11,12^. The CRL4 complex specifically consists of cullin4A (CUL4A) or cullin4B (CUL4B), the RING protein RBX1, the adaptor protein DDB1 (damaged DNA binding 1), and one of several substrate receptors, including CRBN (Fig. 1A)^9,13,14^. CRLs are regulated by dynamic modification of the cullin subunit with the ubiquitin-like modifier Nedd8^11,12,15,16^. Like Ub, Nedd8 requires the sequential action of Nedd8-specific E1, E2 and E3 enzymes. Nedd8-modification of cullins is required for E3 ligase activity, by exposing RBX1 and thereby positioning the associated E2 enzyme in proximity of substrates^16^. Conversely, deneddylation of cullins is required for the exchange of the substrate receptor module^17,18^. Cullin deneddylation is mediated by the multisubunit COP9 signalosome (CSN) ^19,20^. Thus, cullin Nedd8-modification is highly dynamic, with neddylation required for E3 ligase activity and deneddylation required for substrate receptor exchange.

**Figure 1.**
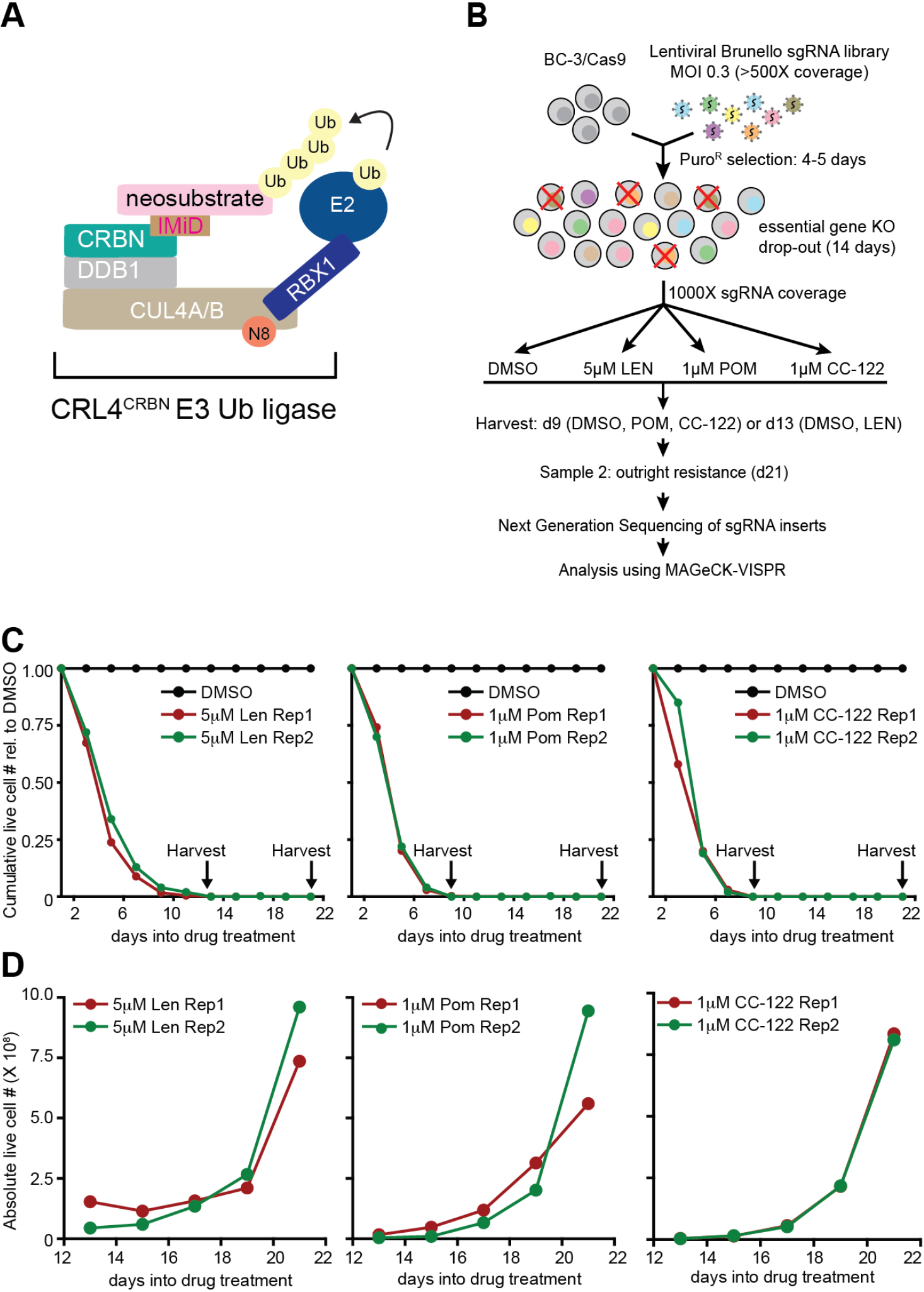
CRISPR-Cas9 inactivation based CM resistance screens in the PEL cell line BC-3. (A) Schematic of the CM-bound CRL4^CRBN^ E3 Ubiquitin ligase complex. (B) Experimental outline of CM resistance screens. (C) Cumulative live cell counts of BC-3/Cas9 over the duration of the screens. Cell numbers were normalized to those from DMSO treated control cells. Arrows indicate early and late time points at which a subset of cells were taken from the pool for analysis of sgRNA distribution. (D) Absolute live cell counts of LEN, POM or CC-122 treated BC-3/Cas9 cells at late time points in the screen shows emergence of proliferating cell pools under continued drug treatment.

CMs bind to CRBN within the substrate recognition surface and thereby enable acquisition of neosubstrates, resulting in their polyubiquitination and proteasomal degradation^21–26^. In multiple myeloma, CM-induced degradation of neosubstrates IKZF1 and IKZF3 and the subsequent downregulation of their transcriptional target IRF4 is thought be a major trigger of CM toxicity^21,22^. Degradation of the neosubstrate CK1α, which is preferentially targeted by LEN, is thought to explain the efficacy of LEN in del(5q) MDS^27^. Recently, CMs have been shown to induce efficient cell death in PEL cell lines^6,7^ and LEN is currently part of a clinical trial in PEL (ClinicalTrials.gov Identifier: NCT02911142). As in multiple myeloma^28^, the transcription factor IRF4 is a highly essential survival gene in PEL cells and its expression is reduced upon treatment with CMs^6,7,29^. Surprisingly, however, in this setting, downregulation of IRF4 and CM toxicity were independent of IKZF1 and/or IKZF3^7^. In PEL cell lines, CK1α is furthermore commonly essential and its degradation represents an IRF4-independent arm of CM toxicity^7^. The individual or combined re-expression of CK1α and IRF4 significantly, but incompletely, rescued PEL cell lines from CM-induced cell death, suggesting that additional mechanisms of CM action exist in PEL. Thus, further mechanistic investigation of CM toxicity in this and other cell types is warranted.

Genome-wide CRISPR screens represent a powerful tool to probe for genes whose inactivation confers drug resistance^30,31^. In order to identify CM effectors, we applied this unbiased screening approach to identify genes whose inactivation confers resistance to LEN, POM, or CC-122 in PEL cells. Our results suggest that decreased CRL4^CRBN^ activity is the dominant pathway to resistance and identify several novel regulators of CRL4^CRBN^. We validate the E2 enzyme UBE2G1 and the Nedd8-specific protease SENP8 as genes whose inactivation can confer resistance to LEN and/or POM. Inactivation of SENP8 indirectly increased the levels of unneddylated CUL4A/B, which interferes with LEN toxicity in a dominant negative manner.

## Methods

### Cell lines and reagents

All parental cell lines were maintained as described recently^29^. Lenalidomide was from Cayman Chemicals (Ann Arbor, MI) and pomalidomide and CC-122 were from Selleck Chemicals (Houston, TX). All compounds were dissolved in DMSO (Sigma-Aldrich, St. Louis, MO). Lentiviruses were prepared as described previously and functionally titrated^29^. Generation and validation of clonal Cas9-expressing BC-3 and BCBL-1 cells were described^29^. Clonal BC-3 CRBN KO cells were described previously^7^. The lentiviral Brunello CRISPR KO library was obtained from Addgene (Cambridge, MA)^32^.

### Genome-wide CRISPR screens

Clonal Cas9 expressing BC-3 cells were transduced with the Brunello library at ~500X coverage and an MOI of ~0.3 and selected with puromycin as described^29^. Cells were passaged for two weeks to allow drop-out of sgRNAs targeting essential genes from the cell pool^29^. Then, cells were treated with 5μM LEN, 1μM POM, 1μM CC-122 or DMSO, in two replicates per condition. Cells were passaged every 2-3 days, while maintaining sgRNA coverage. Samples corresponding to 1000X sgRNA coverage were harvested on day 9 into POM/CC-122/DMSO or day 13 into LEN/DMSO treatment. A second sample for all conditions was taken on day 21. Preparation of genomic DNA and next generation sequencing libraries was performed as described^29^. Trimmed reads were aligned to the Brunello sgRNA library^32^ using Bowtie and raw read counts were generated for each sgRNA as previously described^29^ (Table S1). To reduce statistical noise and false positives, sgRNAs with read counts in the lowest 5% in the DMSO control were excluded from further analyses (Table S2). These sgRNAs include those with few copies in the library and/or those that target highly essential genes. Filtered read counts were analyzed with MAGeCK-VISPR^33^ to test for enrichment of genes in treatment groups using total read normalization. sgRNA and gene level results for enrichment are in Tables S3 and S4, respectively. Raw data are available under GEO accession number <u>GSE122040</u>, using reviewer token ixexicugzfkbzul).

### Cloning

Lentiviral sgRNA vectors were based on pLenti-guide puro^30^ (Addgene#52963), see Table S5 for sgRNA sequences and primers. For lentiviral protein expression, cDNA sequences were placed into pLC/P2Ahygro (CRBN, UBE2G1, SENP8) or pLC/P2Apuro (CUL4A WT/K705R). Detailed cloning procedures are described in the supplemental methods. Primer sequences are listed in Table S6 and cDNA nucleotide sequences are listed in Table S7.

### Generation of cell pools by lentiviral transduction

For KO cell pools prepared for this study (CUL4A, CUL4B, UBE2G1, ILF3, YPEL5, SENP8) clonal Cas9 expressing BC-3 cells were transduced with lentiviral sgRNA vectors based on pLenti-guide puro (Addgene #52963)^34^, at an MOI of 1.5. For lentiviral protein expression, cells were transduced at an MOI <1, media were changed 24 hours after transduction, and cells were selected with hygromycin or puromycin for 5 or 3 days, respectively. Western Blot analyses were used to confirm gene inactivation or expression of the target proteins.

### Cell counting, Western Blotting, and IC_50_ dose-response assays

IC50 assays were performed as described^7^. For growth curves, 4×10^5^ cells were treated with the indicated concentrations of LEN, POM, CC-122 or DMSO and live cell numbers were measured every 2 days using the Cell Titer Glo 2.0 kit (Promega, Madison, WI). At each time point, resulting data were used to calculate absolute live cell numbers, based on manual counting of a subset of samples, and cell concentrations were adjusted to 2 or 3 × 10^5^ cells/ml. Cells were pelleted and resuspended in fresh media containing drug/DMSO at each passage. At each time point, live cell numbers of the drug treated samples were normalized to their respective DMSO samples. Matched aliquots were washed twice with PBS and processed for quantitative Western Blot analysis ((LI-COR) as reported^7,29^. Settings for primary antibodies are listed in Table S8.

## Results

### Genome-wide CRISPR/Cas9 based positive selection screens against CMs in PEL

We conducted genome-wide CRISPR/Cas9 resistance screens against LEN, POM and CC-122 in the commonly used PEL cell line BC-3 (Fig.1B) to identify effectors of CMs in an unbiased approach. Half maximal inhibitory concentrations (IC_50_) of these drugs in BC-3 are ~2μM for LEN, 213 nM for POM, and 117 nM for CC-122 (Fig. S1). Cas9-expressing BC-3 cells were transduced with the Brunello genome-wide sgRNA library and cultured for two weeks following antibiotic selection to allow for depletion of cells with sgRNAs targeting essential genes^29^. The resulting cell pool was divided and treated with DMSO or lethal concentrations of CMs, i.e. 5μM LEN, 1μM POM, or 1μM CC-122. High concentrations were chosen to assay for robust CM resistance under stringent conditions. A first sample was collected when a minimum of live cells remained: on day 14 for LEN, or day 9 for POM or CC-122 (Fig. 1C). A second sample was harvested after a proliferating cell pool emerged under drug treatment (Fig. 1D). The later time point was intended to assay for complete resistance, while the early time was included to detect more subtle delays in CM toxicity. sgRNA composition over two replicate screens per drug was assessed by next generation sequencing and analyzed with the MAGeCK algorithm^33,35^.

For each drug, sgRNAs targeting fewer genes were significantly enriched in the live cell population at late time points than at early time points (Fig. 2A-F). The number of hits furthermore decreased with increasing toxicity of the CM, such that only inactivation of CRBN appeared to confer complete resistance to CC-122 (Fig. 2F). An enrichment for guides against CRBN was expected, because its inactivation has previously been shown to confer CM resistance in PEL^6,7^ and MM cell lines^21,22^. Other genes with strong sgRNA enrichments in at least a subset of settings include: the CRL4 cullin subunit CUL4B; GLMN, a previously identified binding partner of RBX1^36^; and the E2 Ub conjugating enzyme UBE2G1, which has very recently been implicated as the preferred E2 enzyme for ubiquitin chain elongation by CRL4^37,38^. sgRNAs targeting four other genes (ILF2, ILF3, YPEL5, SENP8) were highly and specifically enriched in LEN-resistant cells (Table S4, Fig. 2G), and therefore represent novel candidates for modulators of LEN toxicity. Of these, ILF2 and ILF3 have previously been identified as candidate interacting partners of several CRL complexes^39^, but have not been functionally investigated as potential regulators of CRL activity and also serve unrelated functions. YPEL5 has been identified as a candidate binding partner of the COP9 signalosome subunit COPS5^40^, but also participates in an unrelated E3 Ub ligase complex^41^, among other roles. SENP8 (also known as DEN1 or NEDP1) encodes a Nedd8-specific cysteine protease that has recently been shown to remove Nedd8 modifications from proteins other than cullins, specifically the Nedd8 conjugation machinery, including Nedd8-specific E1 and E2 enzymes^42^. While SENP8-inactivation has recently been shown to cause decreased neddylation of CUL1 and CUL5^42^, SENP8 has not previously been implicated in the regulation of CRL4 activity or CM toxicity.

**Figure 2.**
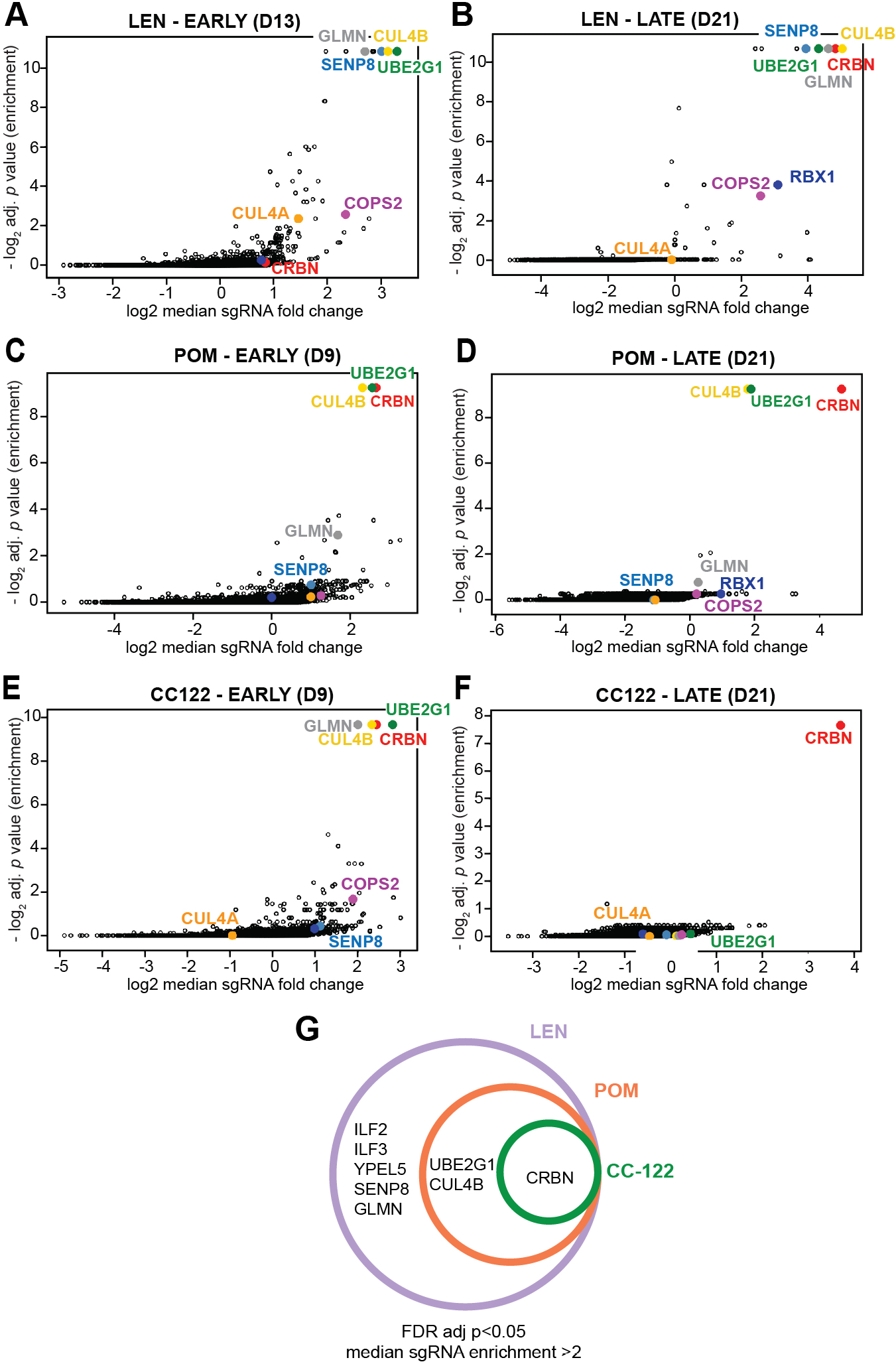
CRISPR screens identify candidates for genes whose inactivation confers CM resistance. (A-F) Gene level analysis of changes in sgRNA distribution in cell pools harvested at early time points (day 13 for LEN or day 9 for POM and CC-122, panels A, C, E) or late time points (day 21, panels B, D, F). The most relevant genes are highlighted and labeled using identical colors in all panels. (G) Venn diagram depicting overlap of most confident candidates for genes whose inactivation confers outright resistance against LEN, POM or CC-122. Cutoffs used were FDR adj. *p* <0.05 and a median sgRNA fold change > 2 on day 21.

Both the experimental settings for our screens and cut-offs for data analyses presented in Fig. 2 were selected for high stringency. Lowering statistical cutoffs identifies additional genes as candidates for modulators of CM toxicity, among known components or regulators of CRL4, such as RBX1 and CSN subunits (see Tables S3-4). We note that DDB1, RBX1, and several CSN subunits are potentially essential in PEL based on our reported essentiality screens^29^ (Fig. S2), which would prevent enrichments of sgRNAs targeting these genes in the CM resistant cell population. Based on the prominent enrichments for CUL4B, UBE2G1, ILF2/3, YPEL5 and SENP8-specific sgRNAs, these genes were selected for further validation.

### Inactivation of CUL4A or CUL4B confers resistance to lenalidomide and delays toxicity of pomalidomide

Our detection of enrichments for CUL4B sgRNAs is interesting, because it suggests that overall CUL4A and CUL4B expression levels in BC-3 cells is limiting for the efficacy of even high concentrations of CMs. To confirm this result, we transduced Cas9 expressing BC-3 cells with lentiviruses carrying two independent sgRNAs each targeting CUL4B or CUL4A, or an established negative control guide targeting the non-coding locus AAVS1. A previously reported CRBN knockout BC-3 clone served as a positive control for CM resistance^7^. Indeed, individual CRISPR/Cas9-mediated inactivation of CUL4B substantially delayed the toxicity of POM and conferred robust resistance to LEN (Fig. S3A-C). Inactivation of CUL4A similarly conferred significant resistance to LEN, although CUL4A-specific sgRNAs were not enriched in our screens (Fig. S3D-E). Inactivation of CUL4A had a smaller effect on the response to POM (Fig. S3F). Overall, these data confirm that CUL4B and CUL4A are both expressed in PEL and contribute to CRL4^CRBN^-dependent CM toxicity in BC-3. The failure for CUL4A sgRNAs to meet our statistical cutoff (Fig. 2) may reflect a relative inefficiency of these guides in the pooled approach and/or, possibly, the high stringency of the experimental setup and cutoffs.

### Inactivation of UBE2G1 confers resistance to lenalidomide and pomalidomide

Remarkably, UBE2G1 was the only E2 Ub conjugating enzyme out of 40 human E2 enzymes with enriched sgRNAs at early (LEN, POM, CC-122) and late (LEN, POM) time points (Figs. 2, S4). To test the hypothesis that inactivation of UBE2G1 confers CM resistance, we infected Cas9 expressing BC-3 cells with lentiviruses carrying two independent sgRNAs targeting UBE2G1 or the sgAAVS1 negative control guide. Inactivation of UBE2G1 was confirmed by Western blot analysis (Fig. 3A). Inactivation of UBE2G1 indeed conferred outright resistance against either LEN or POM (Fig. 3B, C). Although live cell numbers of UBE2G1 knockout cells under CM treatment were lower than those of CBRN-KO cells, these cells continued to proliferate under drug treatment. Re-expression of an sgRNA-resistant UBE2G1 cDNA in the context of UBE2G1 inactivation restored sensitivity to CMs, confirming the specificity of sgUBE2G1-mediated CM resistance (Fig. 3D, E). Western blot analyses of UBE2G1-inactivated cells demonstrate substantial impairment of CM-induced neosubstrate degradation in LEN or POM-treated UBE2G1-inactivated cell pools for IKZF1, IKZF3 and CK1α (Fig. 3F, G), which were analyzed as indicators of overall CRL4^CRBN^ activity. The LEN/POM-induced indirect downregulation of IRF4 was similarly rescued in UBE2G1-inactivated cells compared to the negative control cells. The rescue of LEN and POM-induced phenotypes strongly suggest that UBE2G1 acts at the level of the CM-bound CRL4^CRBN^ complex, most likely as the preferred E2 ubiquitin conjugation enzyme participating in the polyubiquitination of CM neosubstrates in PEL. Validation experiments furthermore confirmed that inactivation of UBE2G1 modestly delays toxicity of the more potent CC-122 but does not confer resistance against this drug (Fig. 3H, I). Thus, other E2 enzymes must be able to compensate for loss of UBE2G1, albeit at lower efficiency. These results parallel two recent reports^37,43^, which similarly identified UBE2G1 as hits in genome-wide CRISPR/Cas9 screens for LEN or POM resistance in multiple myeloma. UBE2G1 was also identified as the preferred E2 in CM-induced CRL4^CRBN^ activity in an E2 enzyme focused CRISPR screen in multiple myeloma^38^. Sievers et al.^37^ and Lu et al.^38^ confirmed a role for UBE2G1 in CM-induced toxicity in MM and proposed a “prime-extend” mechanism, whereby the E2 enzyme UBE2D3 mediates substrate mono-ubiquitination, while UBE2G1 extends the Ub chain via addition of further K48-linked Ub moieties. Inactivation of UBE2D3 alone, which did not score in our screens, did not confer significant CM resistance in PEL, suggesting the existence of redundant E2 enzymes that can fulfill this function in our system (Fig. S5). Overall, our data suggest a preference for CRL4^CRBN^ for UBE2G1, similar to that seen in multiple myeloma^37,38,43^.

**Figure 3.**
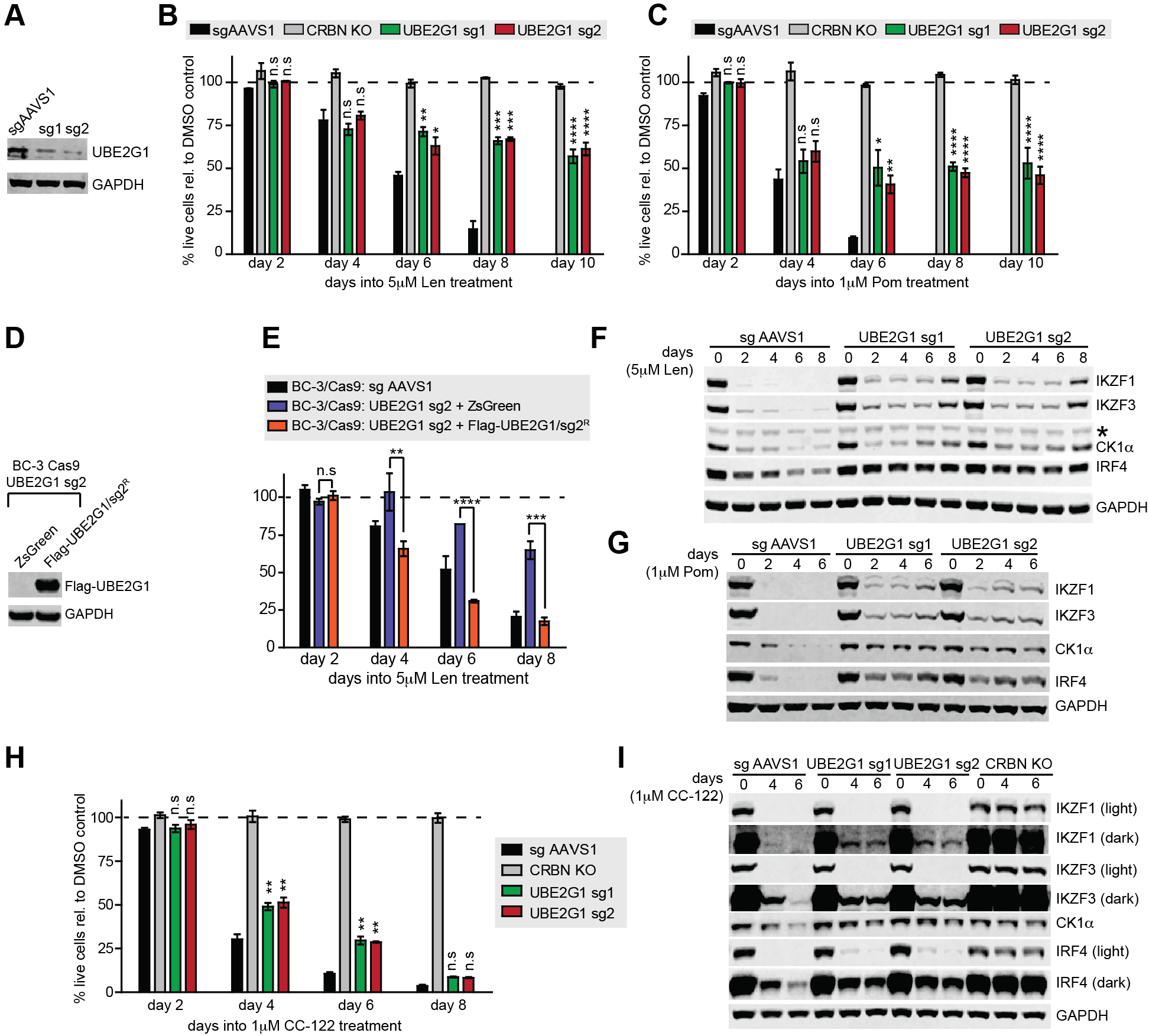
Inactivation of UBE2G1 confers resistance to LEN and POM, but not CC-122. (A) Representative Western blot analysis confirms efficient CRISPR-induced inactivation of UBE2G1 in BC-3/Cas9 cells by two independent UBE2G1-specific sgRNAs (sg1, sg2). sgAAVS1 is a negative control guide. GAPDH served as loading control. (B-C) Growth curve analyses of UBE2G1 inactivated BC-3/Cas9 cells after treatment with 5μM LEN (B) or 1μM POM (C). BC-3/Cas9 cells transduced with sgAAVS1 were included as a CM-sensitive negative control, while previously described clonal CRBN-inactivated BC-3 cells (CRBN KO) served as a positive control for complete CM resistance. Absolute live cell numbers were normalized to corresponding DMSO vehicle treated cells at each passage (represented by the dotted line). These experiments were done in parallel with SENP8 inactivated cells shown in Figures 4B and 4C and thus share common negative and positive controls for several or all replicates. (n=3, error bars represent SEM). (D) Representative Western blot analysis (anti-Flag) confirms lentiviral expression of sg2-resistant, Flag-tagged UBE2G1 in UBE2G1-inactivated BC-3/Cas9 cells (UBE2G1 guide RNA sg2). GAPDH served as loading control. (E) Growth curve analysis of UBE2G1-inactivated BC-3/Cas9 cells (UBE2G1 guide RNA sg2) expressing Flag tagged UBE2G1 (sg2-resistant; UBE2G1/sg2^R^) or control ZsGreen following treatment with 5μM LEN confirms restoration of CM sensitivity upon UBE2G1 re-expression. BC3/Cas9/sg AAVS1 were included as a LEN-sensitive positive control. Live cell counts were normalized to corresponding DMSO vehicle treated cells, which are represented by the dotted line. These assays were run in parallel with those in Figure 4E and thus share common controls (n=3, error bars represent SEM). (F-G) Representative time course Western blot analyses of CM neosubstrates IKZF1, IKZF3, and CK1α and IRF4 in WT or UBE2G1-inactivated BC-3/Cas9 cells upon treatment with 5μM LEN (F) or 1μM POM (G). Lysates are matched with growth curves shown in panels B and C, respectively. GAPDH served as loading control. The asterisk marks a non-specific band. (H) Growth curve analyses of UBE2G1-inactivated BC-3/Cas9 cells treated with 1μM CC-122. sgAAVS1 and CRBN KO cells were included as negative and positive controls, respectively Live cell counts were normalized to corresponding DMSO vehicle treated cells, represented by the dotted line. (n=3, error bars represent SEM). (I) Representative Western blot analysis of IKZF1, IKZF3, CK1α and IRF4 at indicated time points confirms a delay in neosubstrate degradation and IRF4 downregulation in UBE2G1-inactivated BC-3/Cas9 cells compared to control pools as in panel H. GAPDH served as loading control. Statistical analyses throughout this figure were done by unpaired *t*-tests comparing specified conditions to corresponding sgAAVS1 or ZsGreen controls. * *p*<0.05, ** *p*<0.01, *** *p*<0.001, **** *p*<0.0001, n.s. not significant.

### Inactivation of SENP8 specifically confers resistance to lenalidomide

SENP8, ILF2, ILF3, and YPEL5 scored specifically for LEN but not POM or CC-122. To test effects of their inactivation on CM toxicity, we targeted each gene using two sgRNAs. Except for ILF2, where efficient inactivation was not achieved (not shown), Western blot analyses confirmed robust target inactivation (Fig. S6A, C, 4A). Inactivation of ILF3 and YPEL5 modestly delayed LEN toxicity (Fig. S6B, D), validating ILF3 and YPEL5 as modulators of CM toxicity. In contrast, inactivation of SENP8 conferred pronounced resistance to LEN (Fig. 4B). Since SENP8 has not been implicated in CRL4 function or CM resistance to date, we decided to further investigate the role of SENP8. As predicted by the screens, inactivation of SENP8 delayed and attenuated toxicity of POM, but did not confer resistance at later time points into drug treatment (Fig. 4C). Observed effects were specific to SENP8 inactivation, since re-expression of sgRNA resistant SENP8 restored LEN-sensitivity (Fig. 4D, E). Western blot analyses of SENP8-inactivated BC-3 cells demonstrate rescue of IRF4 and the neosubstrates IKZF1, IKZF3 and CK1α upon treatment with LEN but not with POM (Fig 4F, G). Similar results were obtained in a second PEL cell line, BCBL-1, where inactivation of SENP8 also conferred robust resistance to LEN and delayed the toxicity of POM (Fig. 4H-J). Overall, these results show that inactivation of SENP8 renders cells less susceptible to LEN toxicity.

**Figure 4.**
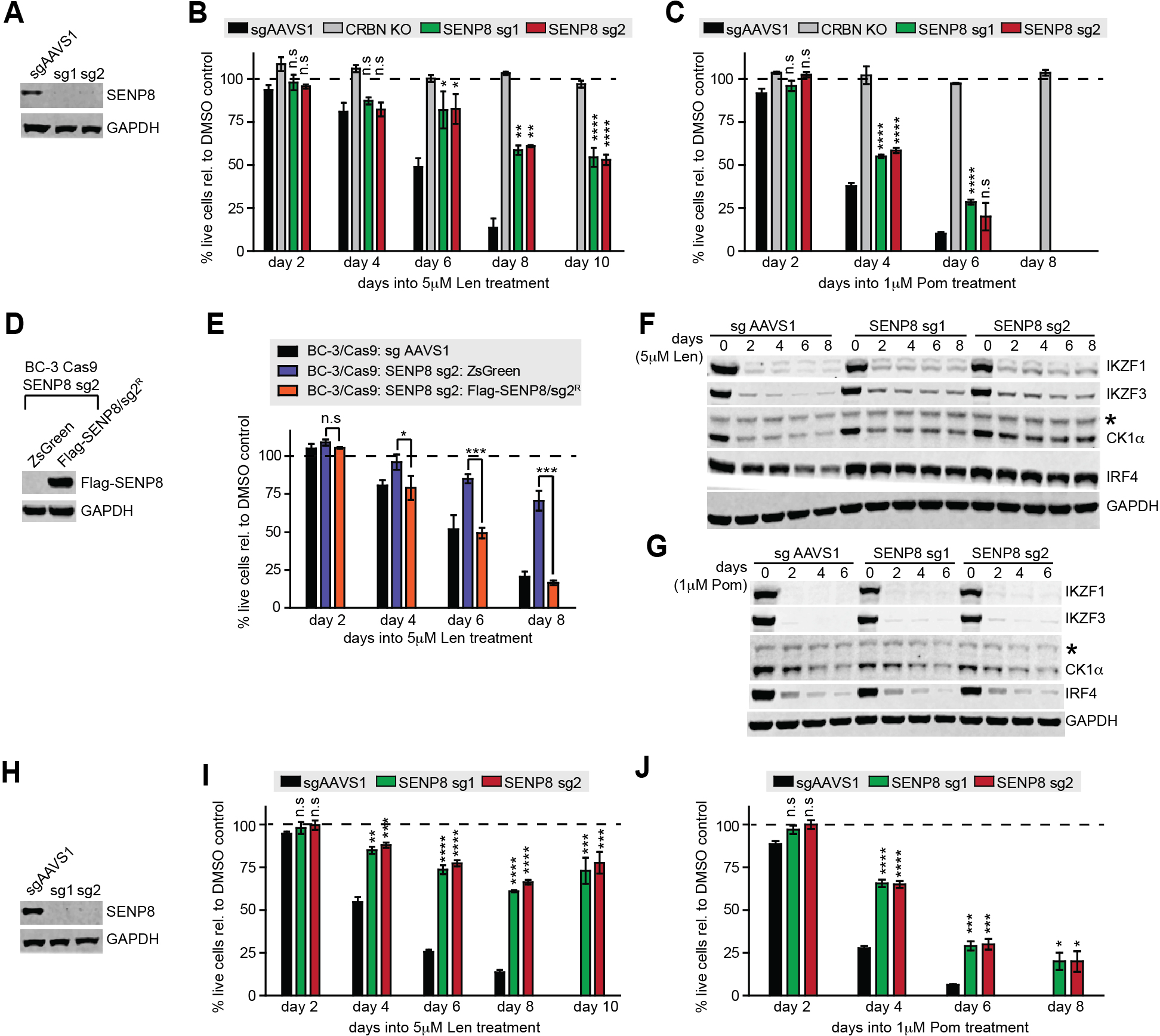
Inactivation of SENP8 confers resistance to LEN but not POM. (A) Representative Western blot analysis confirms efficient CRISPR-induced inactivation of SENP8 in BC-3/Cas9 cells by two independent SENP8-specific sgRNAs (sg1, sg2). sgAAVS1 served as negative control guide, GAPDH served as loading control. (B-C) Growth curve analyses of SENP8-inactivated BC-3/Cas9 cells after treatment with 5μM LEN (B) or 1μM POM (C). BC-3/Cas9 cells transduced with sgAAVS1 were included as a CM-sensitive negative control, while previously described clonal CRBN-inactivated BC-3 cells (CRBN KO) served as a positive control for complete CM resistance. Absolute live cell numbers were normalized to corresponding DMSO vehicle treated cells at each passage (represented by the dotted line). These experiments were done in parallel with UBE2G1-inactivated cells shown in Figures 3B and 3C and thus share common negative and positive controls for several or all replicates. (n=3, error bars represent SEM). (D) Representative Western blot analysis (anti-Flag) confirms lentiviral expression of sg2-resistant, Flag-tagged SENP8 in SENP8-inactivated BC-3/Cas9 cells (SENP8-specific guide RNA sg2). GAPDH served as loading control. (E) Growth curve analysis of SENP8-inactivated BC-3/Cas9 cells (SENP8 guide RNA sg2) expressing Flag tagged SENP8 (SENP8/sg2^R^) or control ZsGreen following treatment with 5μM LEN confirms restoration of LEN sensitivity upon SENP8 re-expression. BC3/Cas9/sg AAVS1 were included as a LEN-sensitive positive control. Live cell counts were normalized to corresponding DMSO vehicle treated cells, which are represented by the dotted line. These assays were run in parallel with those in Figure 3E and thus share a common sg AAVS1 control (n=3, error bars represent SEM). (F-G) Representative Western blot analyses of IKZF1, IKZF3, CK1α and IRF4 at indicated time points confirm incomplete neosubstrate degradation and IRF4 downregulation in SENP8-inactivated BC-3/Cas9 cells compared to sgAAVS1 transduced BC-3/Cas9 control cells upon treatment with 5μM LEN (F) or 1μM POM (G). Lysates are matched with growth curves shown in panels B and C, respectively. GAPDH served as loading control. The asterisk marks a non-specific band. (H) Representative Western blot analysis confirms efficient CRISPR-induced inactivation of SENP8 in BCBL-1/Cas9 cells by two independent SENP8-specific sgRNAs (sg1, sg2). sgAAVS1 served as negative control guide, GAPDH served as loading control. (I-J) Growth curve analyses of SENP8-inactivated BCBL-1/Cas9 cells after treatment with 5μM LEN (I) or 1μM POM (J). sgAAVS1 transduced cells served as a negative control. Live cell counts were normalized to corresponding DMSO vehicle treated cells represented by the dotted line (n=3, error bars represent SEM). Statistical analyses throughout this figure were done by unpaired *t*-tests comparing specified conditions to corresponding sgAAVS1 or ZsGreen controls. * *p*<0.05, ** *p*<0.01, *** *p*<0.001, **** *p*<0.0001, n.s. not significant.

### SENP8 inactivation results in accumulation of unneddylated CUL4A/B

A recent report showed that SENP8 cleaves Nedd8 from non-cullin neddylation substrates, including the Nedd8-specific E2 enzyme UBE2M and other components of the neddylation machinery^42^. Accumulation of aberrant neddylated forms of these proteins appears to render neddylation of cullins in SENP8 inactivated cells inefficient, as shown for CUL1 and CUL5 in HeLa and HEK293T cells^42^. The same study reported no effect on the neddylation status of CUL4, however. To begin to address the role of SENP8 in PEL, we probed WT and SENP8-inactivated BC-3 cells for Nedd8. As reported for 293T^42^, SENP8-inactivated BC-3 showed a depletion of unconjugated Nedd8 and an accumulation of several Nedd8-modified proteins, presumably substrates for Nedd8 deconjugation by SENP8. In both BC-3 and BCBL-1 cells, there was an increase in the levels of unneddylated CUL4A and CUL4B (Fig. 5B-C). Importantly, the expression of CRBN was unchanged in SENP8-inactivated cell pools (Fig. 5B). Therefore, the effect of SENP8 on CM efficacy is most likely at the level of reduced CUL4A/B neddylation. CUL4A/B neddylation and LEN-sensitivity in the context of SENP8 inactivation could not be restored by overexpression of either active or precursor Nedd8 (Fig. S7). These data suggest that the accumulation of unneddylated CUL4A/B is not a consequence of limiting pools of unconjugated Nedd8.

**Figure 5.**
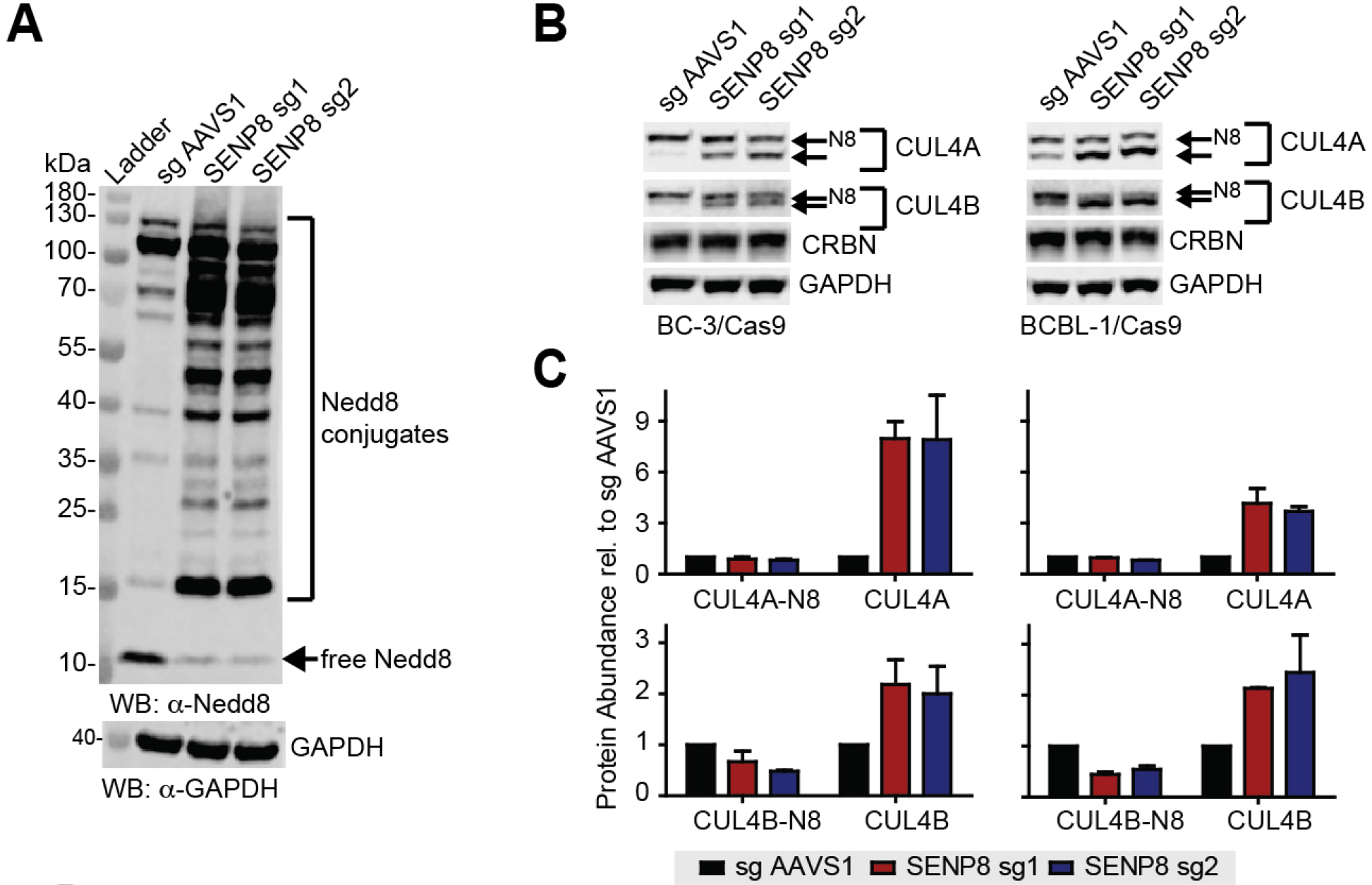
SENP8 inactivation leads to an increase of unneddylated CUL4A/B and depletion of free Nedd8. (A) Representative Western blot analysis of SENP8-inactivated and control sgAAVS1-transduced BC-3/Cas9 cells indicates a strong depletion of unconjugated Nedd8 and an increase in Nedd8 conjugated proteins. GAPDH served as loading control. SENP8 sg1 and sg2 are two independent SENP8-specific sgRNAs used. (B) Representative Western blot analyses of CUL4A, CUL4B and CRBN in SENP8 inactivated BC-3/Cas9 and BCBL-1/Cas9 cells indicate an increase in unneddylated CUL4A/B and no significant change in CRBN protein levels. Neddylated (N8) and unneddylated forms of CUL4A/B are indicated by arrows. GAPDH served as loading control. (C) Quantitative analysis of the data shown in panel B over replicates. Protein levels were first normalized to corresponding GAPDH levels and then to sg AAVS1 levels (n=2, error bars represent SEM).

### CRBN overexpression restores sensitivity to lenalidomide in the context of SENP8 inactivation

Because expression of Nedd8-modified CUL4A/B was not strongly reduced in the context of SENP8 inactivation, we reasoned that unneddylated CUL4A/B might dominant negatively interfere with CM toxicity. Interference could be through sequestration of CRBN on unneddylated complexes. Alternatively, increased substrate receptor exchange facilitated by CUL4 deneddylation could cause competition with alternative substrate adaptors. In either case, overexpression of CRBN would be expected to drive increased association of CRBN /DDB1 with active CRL4 complexes under conditions where activation of CUL4 by neddylation is limiting due to loss of SENP8. Indeed, lentiviral overexpression of Flag-tagged CRBN restored full sensitivity of BC-3 cells to LEN in the context of SENP8 inactivation (Fig 6A). Western blot analysis confirmed restoration of LEN-induced neosubstrate degradation and IRF4 downregulation in CRBN overexpressing cells (Fig 6B). We finally directly tested whether overexpression of non-neddylatable CUL4A K705R mutant dominant negatively interferes with CM toxicity (Fig. 6C). While overexpression of WT CUL4A did not affect the toxicity of LEN or POM, K705R-mutant CUL4A expressing BC-3 were completely resistant to high concentrations of either LEN or POM, which formally shows that unneddylated CUL4 can drive resistance to CMs in a dominant negative manner (Fig 6D-E).

**Figure 6.**
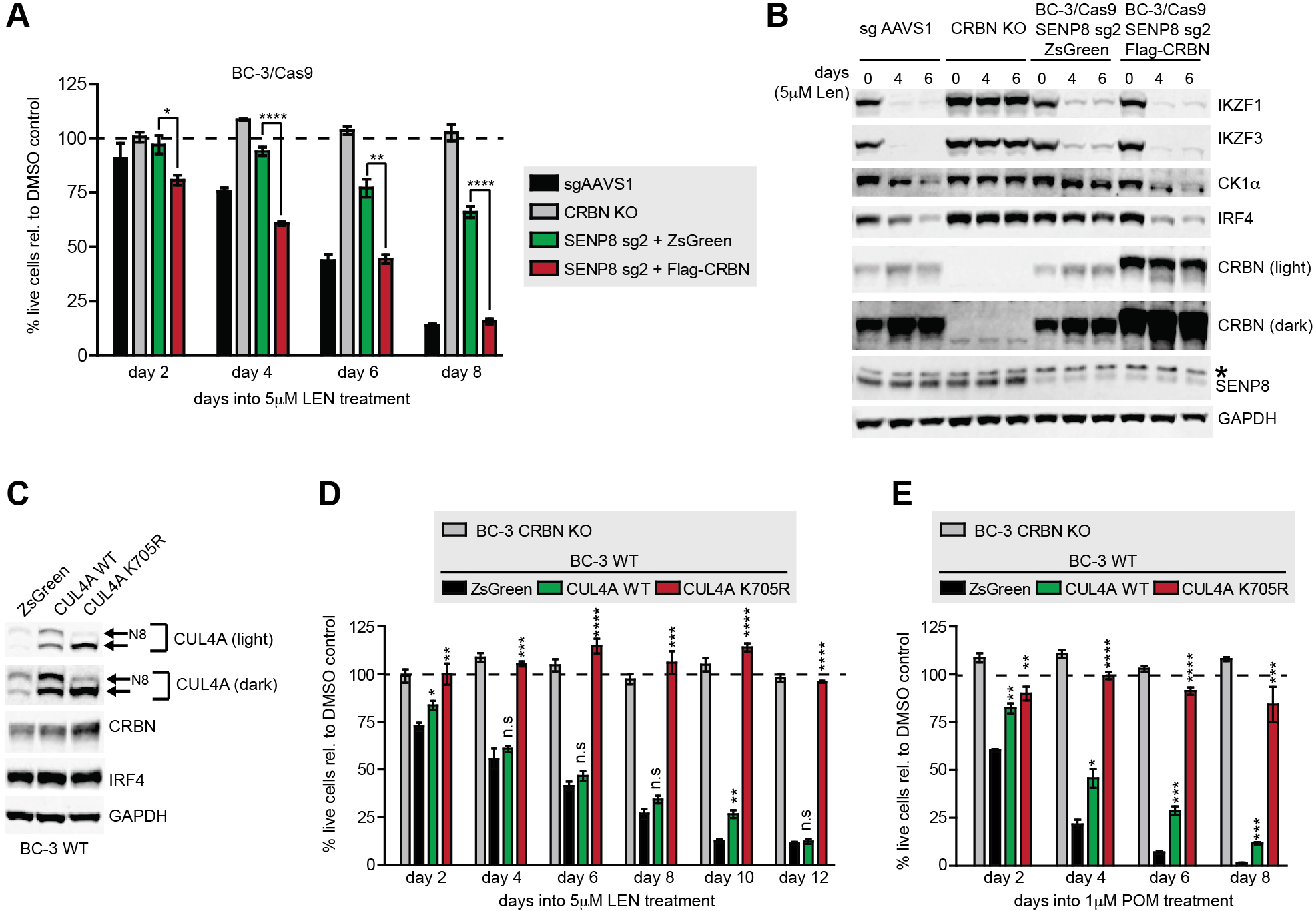
Overexpression of CRBN restores LEN sensitivity in SENP8-inactivated cells. (A) Growth curve analyses of SENP8-inactivated BC-3/Cas9 cells (using SENP8-specific sg2) lentivirally transduced to overexpress Flag-tagged CRBN or ZsGreen and treated with 5μM LEN. sgAAVS1-transduced cells serve as a control for LEN sensitive BC-3 and CRBN inactivated BC-3 cells (CRBN KO) were used as a control for LEN resistant cells. Live cell counts were normalized to DMSO vehicle treated cells, represented by the dotted line (n=3, error bars represent SEM). (B) Representative Western blot analysis of IKZF1, IKZF3, CK1α, IRF4, CRBN and SENP8 at indicated time points into the experiment shown in panel A. GAPDH served as loading control. (C) Representative western blot analysis of BC-3 cells transduced with lentiviral vectors constitutively expressing wild-type (WT) CUL4A, K705R mutant CUL4A, or ZsGreen. Membranes were probed for CUL4A, CRBN, IRF4 or the loading control GAPDH. Neddylated (N8) and unneddylated forms of CUL4A are indicated by arrows. (D-E) Growth curve analyses of BC-3 cells from panel C and treated with 5μM LEN (D) or 1μM POM (E). CRBN inactivated BC-3 cells (CRBN KO) were used as a control for LEN resistant cells. Live cell counts were normalized to DMSO treated control cells, as represented by the dotted line (n=3, error bars represent SEM). For Western Blot analysis of this experiment, see Fig. S8. Statistical analyses throughout this figure were done by unpaired *t*-tests comparing specified conditions to corresponding ZsGreen controls. * *p*<0.05, ** *p*<0.01, *** *p*<0.001, **** *p*<0.0001, n.s. not significant.

## Discussion

Here we report genome-wide CRISPR KO screens for genes whose inactivation confers resistance to three generations of CMs, i.e. LEN, POM and CC-122, in the PEL cell line BC-3. Our study confirms that CRBN is strictly required for CM toxicity. In addition, screens and validation experiments show that the E2 Ub conjugating enzyme UBE2G1 and the deneddylase SENP8 are required for optimal CM toxicity in PEL. While our work was in progress, similar screens against LEN^37^ or POM^43^ were reported in multiple myeloma. These screens and an additional E2-targeted CRISPR screen^38^ similarly identified loss of UBE2G1 as a mechanism of LEN and POM resistance in multiple myeloma. Our screens additionally identified SENP8 and other novel candidates as modulators of CM toxicity in PEL.

Our results suggest that genes that confer robust CM resistance are mostly, or possibly exclusively, modulators of CRL4^CBRN^ activity. This can be explained by our previous finding that mechanisms of CM action in PEL are multifactorial^7^. Interestingly, the number of genes whose inactivation confers resistance decreases with increased efficacy of the drug. Even relatively subtle impairments in CRL4^CRBN^ activity, e.g. following loss of SENP8, appear to limit LEN toxicity. In contrast, POM and especially CC-122 remain effective under conditions of suboptimal CRL4^CRBN^ activity, such as reduced expression of overall CUL4 or loss of SENP8. Strikingly, the drug target CRBN was the only gene whose inactivation conferred complete resistance to CC-122 in our system.

Amongst the CRL4^CRBN^ regulators identified in our study was the E2 Ub conjugating enzyme UBE2G1. We confirmed that inactivation of UBE2G1 in PEL cells confers resistance to LEN and to POM, but not CC-122. LEN and POM resistance of UBE2G1-inactivated cells was recently also reported in multiple myeloma by Sievers et al.^37^ and Lu et al. ^38^. These groups proposed a “prime-extend” mechanism, where the E2 conjugating enzyme UBE2D3 mediates neosubstrate mono-ubiquitination, while UBE2G1 adds additional Ub moieties for polyubiquitination. Our results extend this role of UBE2G1 to PEL, where CM toxicity is independent of CM-induced IKZF1/3 degradation^7^. Future CRISPR screens in the context of UBE2D3 and/or UBE2G1 inactivation could be used to identify E2 enzymes that can compensate for these enzymes, especially during treatment with CC-122.

Inactivation of either CUL4A or CUL4B conferred significant resistance to LEN, suggesting that total CUL4A/B levels are likely limiting for CM toxicity in BC-3. Limited CUL4 expression could sensitize cells to alterations in CUL4 neddylation status, such as those seen in SENP8 inactivated cells. It is conceivable that certain tumor types, including PEL, have low CUL4A/B expression or that cells with low CUL4A/B expression exist within tumors for various cancers, which may impact the response to CM treatment and increase the likelihood of relapse.

Our screens and validation experiments identified a role for the Nedd8 specific protease SENP8 in the modulation of LEN toxicity and CRL4 activity. Specifically, inactivation of SENP8 leads to accumulation of deneddylated CUL4, perhaps a consequence of aberrant neddylation and reduced activity of the Nedd8 conjugation machinery^42^. Increased amounts of deneddylated CUL4A/B may lead to the sequestration of DDB1/CRBN on inactive CRL4 complexes, thus limiting CM toxicity in SENP8 inactivated cells. Accordingly, overexpression of CRBN restored sensitivity of SENP8 KO cells to LEN. Conversely, overexpression of non-neddylatable CUL4A dominant negatively interfered with CM toxicity and thereby phenocopied SENP8 knockout.

Finally, our study identified and validated ILF3 and YPEL5 as modulators of CM toxicity. ILF2 and ILF3 were recently identified as candidate binding partners of CRL complexes and their roles should be addressed in future studies^39^. YPEL5 is a component of the unrelated GID/CTLH E3 ligase complex. The early time point for LEN identified GID components WDR26 and MEMA, as well as the cognate E2 enzyme of this complex (UBE2H) as candidates for genes with delayed LEN toxicity, perhaps implicating the GID E3 ligase as a regulator of CM toxicity. However, YPEL5 has also been identified as a candidate interacting partner of the CSN subunit COPS5^40^ and has unrelated functions.

In summary, we propose a model where overall expression levels of CRBN, CUL4A, CUL4B, UBE2G1, and SENP8 dictate sensitivity of CMs in PEL (Fig 7A-C). Sequencing and gene expression studies will be required to identify roles of the expression or mutation of these genes in the response to CMs and the development of CM resistance in patients. The expression and/or mutational status of the genes identified in this study may be developed as biomarkers for whether treatment with a more potent CM is indicated upon relapse from LEN or POM. Moving forward, it will be critical to identify the relevant neosubstrate(s) and mechanisms of CM toxicity in PEL. An improved understanding of the mechanisms of CM toxicity might also enable combination therapy with other drugs.

**Figure 7.**
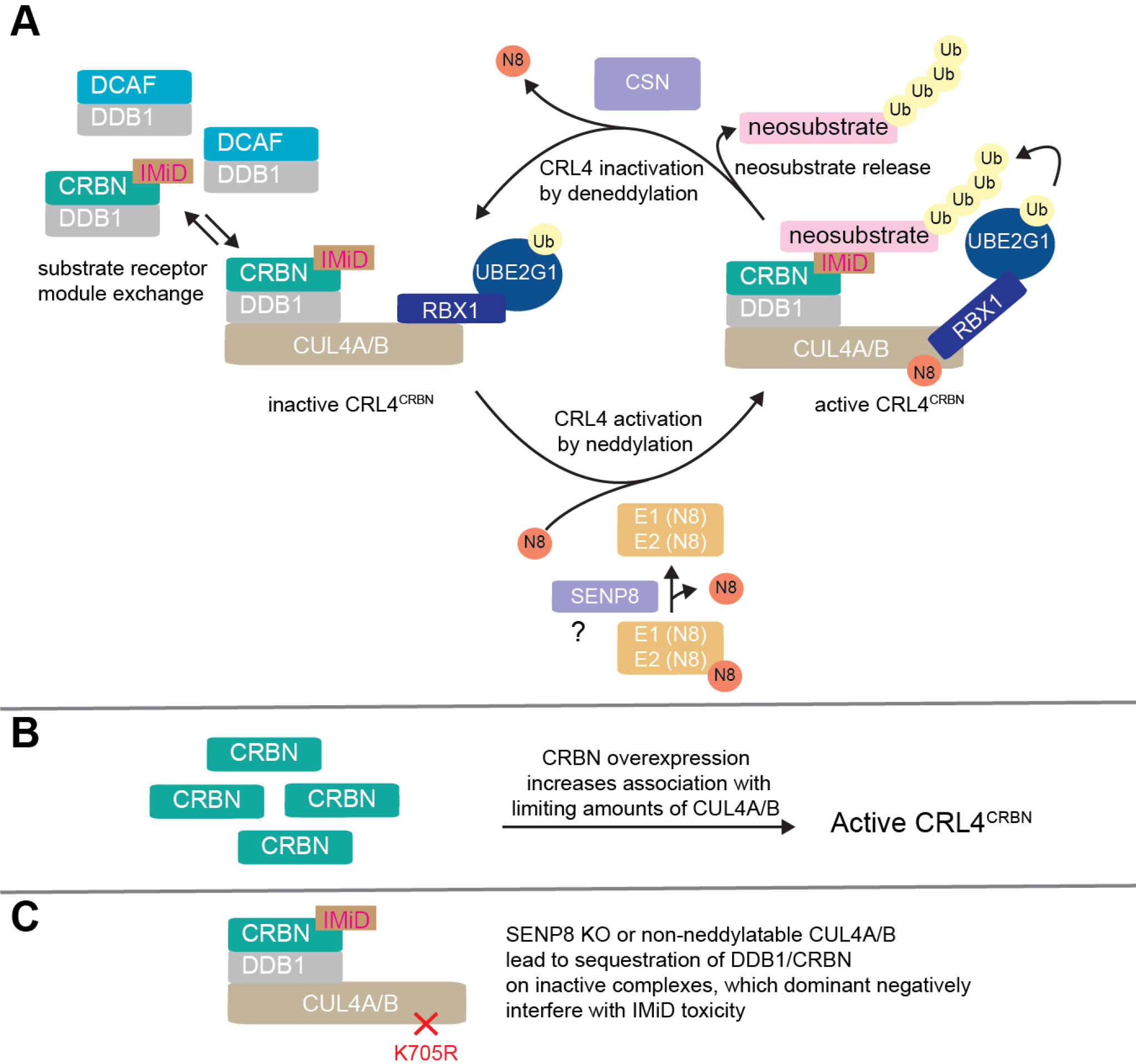
Model of the roles of UBE2G1 and SENP8 in CM toxicity. (A) We hypothesize that UBE2G1 is the preferred E2 ubiquitin conjugating enzyme for polyubiquitination by CM-bound CRL4^CRBN^. SENP8 affects the neddylation status of CUL4A and CUL4B, potentially by cleaving detrimental Nedd8-conjugates from Nedd8-specific E1 and E2 enzymes^42^, and thereby affects CRL4^CRBN^ activity. (B) CRBN overexpression promotes assembly of active CRL4^CRBN^ complexes, likely by outcompeting other DDB1/DCAF substrate recognition modules for limiting pools of CUL4A/B. (C) Unneddylated CUL4A/B, either a consequence of SENP8 inactivation or due to expression of non-neddylatable mutants, interferes with CRL4^CRBN^ activity in a dominant negative manner, potentially by sequestration of DDB1/CRBN on inactive complexes. Our data and this model suggest that CRBN and CUL4A/B expression in BC-3 are limiting. CSN: COP9 signalosome, DCAF: DDB1-CUL4-Associated Factors.

## Acknowledgements

The authors thank the Northwestern University NUSeq Core Facility and the University of Chicago Genomics Facility for their Illumina sequencing services and Kylee Morrison and Samuel Harvey for their feedback on this manuscript. This study was supported by National Institutes of Health, National Cancer Institute Grant R21 CA210904, by Searle and Zell Scholar Awards from the Robert H. Lurie Comprehensive Cancer Center (E.G.), and by Chicago Biomedical Consortium Postdoctoral Award PDR-061 (M.M.). The content is solely the responsibility of the authors and does not necessarily represent the official views of the funding agencies.

## Authorship contributions

A.P. and E.G. designed the study, analyzed data and wrote the manuscript. A.P. conducted all experiments. M.M. prepared a subset of next generation sequencing libraries, analyzed data, and provided feedback on the study and manuscript.

