## Supplementary Methods and Figures for "Genome-wide CRISPR Screens Reveal Genetic Mediators of Cereblon Modulator Toxicity in Primary Effusion Lymphoma"

#### Supplemental Data

- **Supplementary Methods**
- **Supplementary References**
- **Supplementary Figures**

### Genome-wide CRISPR Screens Reveal Genetic Mediators of Cereblon Modulator Toxicity in Primary Effusion Lymphoma

Ajinkya Patil[1], Mark Manzano[1], and Eva Gottwein[1]

[1]Department of Microbiology-Immunology, Feinberg School of Medicine, Northwestern University, Chicago, Illinois, 60611 USA.

**Short Title:** Cereblon Modulator Resistance Screens in PEL

**Corresponding Author:**
Eva Gottwein
Department of Microbiology-Immunology
Northwestern University Feinberg School of Medicine
320 E Superior St., Tarry Bldg., Room 6-735
Chicago, Illinois 60611
USA

**Supplementary Methods**

**Vector design and cloning procedures**

sgRNA sequences and primers are listed in Table S5, all other cloning primers in Table S6, and cDNA nucleotide sequences in Table S7. sgRNAs were designed using the algorithm by Doench et al.[1] and cloned into the BsmBI site of pLenti-guide puro[2] (Addgene# 52963). sgAAVS1 is a published control targeting a non-coding locus[3-5].

Lentiviral expression vectors were based on pLC/ZsGreen-P2A-Hygro and pLC/ZsGreen-P2A-Puro. To construct pLC/ZsGreen-P2A-Hygro, PCR fragments generated using primers 3883/3885 (CMV promoter), 3887/3888 (ZsGreen-P2A), and 3908/3909 (P2A-Hygro$^R$) were Gibson assembled with the large BamHI/EcoRI fragment of pLCE[6] to create a CMV-ZsGreen-P2A-Hygro coding sequence. The ZsGreen-P2A fragment was amplified from pZIP-hCMV-ZsGreen-P2A-Puro-NT4[4], other fragments were from standard cloning vectors. To construct pLC/ZsGreen-P2A-Puro, pLCE-P2A-Puro[4] was digested with BamHI and Gibson assembled with PCR products generated using primers 3883/3885 (CMV promoter) and 3887/3888 (ZsGreen-P2A). This strategy renders the downstream BamHI site unique.

To clone pLC-Flag-UBE2G1/sg2$^R$-P2A-Hygro and pLC-Flag-SENP8/sg2$^R$-P2A-Hygro, pLC/ZsGreen-P2A-Hygro was digested with AgeI and BamHI to remove ZsGreen and Gibson assembled with a PCR product encoding Flag-tagged, sgRNA-resistant UBE2G1 or SENP8. N-terminal Flag tags and silent mutations that disrupt the UBE2G1 sg2 or SENP8 sg2 binding sites were introduced with the PCR primers (Table S6). UBE2G1 cDNA was amplified from Addgene#31407[7] and SENP8 cDNA was amplified from the BC031411 clone obtained from Dharmacon Inc (Catalog# MHS6278-202801152).

To clone pLC/HA-Nedd8-P2A-Hygro and pLC/HA-active Nedd8-P2A-Hygro, pLC/ZsGreen-P2A-Hygro was digested with AgeI and BamHI to remove ZsGreen and subjected to Gibson cloning with a PCR product encoding HA-tagged Nedd8 or HA-tagged active Nedd8. The active Nedd8 construct lacked nucleotides coding for the C- terminal amino acids GGLRQ. The HA tag was added at the N-terminus of both constructs. Nedd8 and active Nedd8 cDNA were amplified from Addgene#18711[8].

To clone pLC/Flag-CRBN-P2A-Hygro, pLC/ZsGreen-P2A-Hygro was digested with AgeI and BamHI to remove ZsGreen and subjected to Gibson cloning with a PCR product encoding Flag tagged CRBN. The Flag tag was added at the N-terminus. CRBN cDNA was amplified from Addgene# 107380[9].

To clone pLC/CUL4A WT-P2A-Puro, pLC/ZsGreen-P2A-Puro was digested with AgeI and BamHI to remove ZsGreen and subjected to Gibson cloning with a PCR product encoding CUL4A. CUL4A cDNA was amplified from the BC008308 clone obtained from Dharmacon Inc (Catalog# MHS6278-202828794). CUL4A K705R construct was generated by overlap PCR using primers 4254 and 4255.

All PCR inserts were verified by Sanger sequencing.

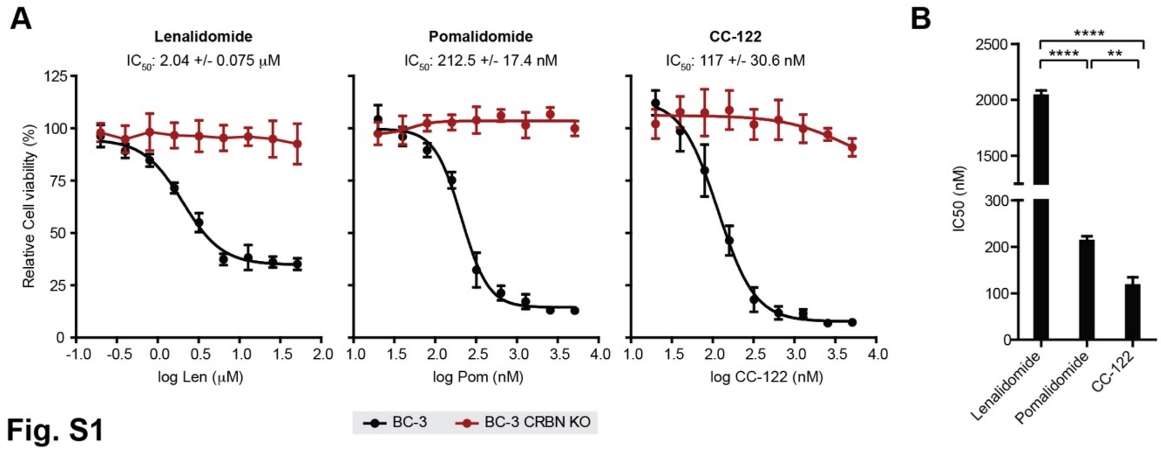


**Figure S1. Increased potency of CMs in BC-3 with later generation. (A)** $IC_{50}$ curves for LEN, POM, and CC-122 in wild-type BC-3 and a previously reported clonal CRBN-inactivated BC-3 cell line (BC-3/CRBN-KO)[4]. (B) $IC_{50}$ values of wild-type BC-3 shown in (A). n=3, error bars represent SEM. Statistical analyses were done by unpaired *t*-tests. ** $p<0.01$, **** $p<0.0001$.

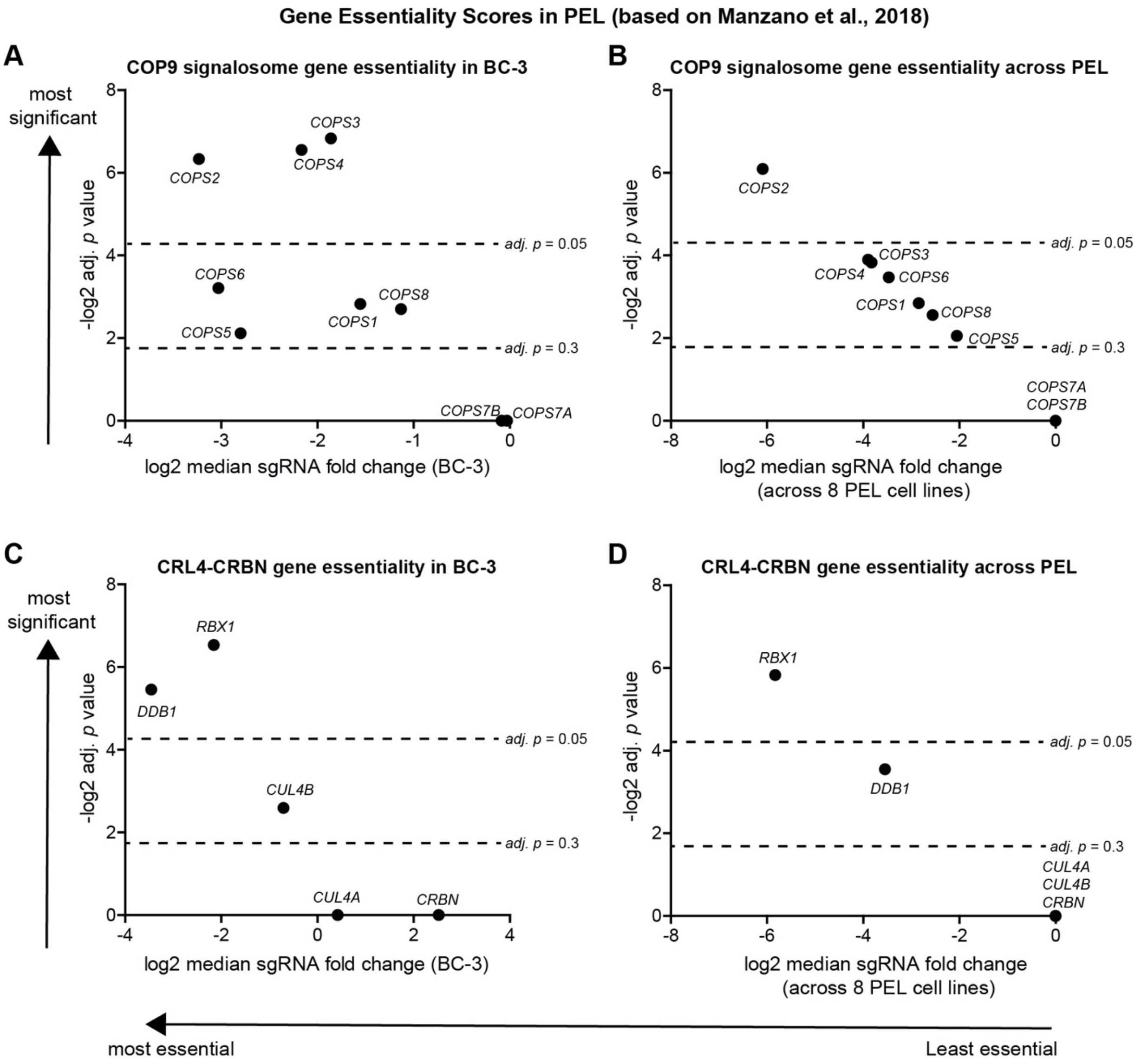


**Figure S2. Essentiality of subunits of the COP9 signalosome and CRL4$^{CRBN}$ for the survival and/or proliferation of PEL cell lines.** (A) Degree of sgRNA depletion for genes encoding CSN subunits in a recently reported CRISPR screen for gene essentiality in BC-3[5]. High -log2 (adj. *p* values) and low log2(median sgRNA fold change) identifies candidates for high essentiality. (B) As in panel (A), except that median values across a panel of 8 distinct PEL cell lines screened in the same study were plotted. (C) as in (A), except that sgRNAs depletion for genes encoding subunits of CRL4$^{CRBN}$ were analyzed in BC-3. (D) as in B, except that sgRNA depletion for genes encoding subunits of CRL4$^{CRBN}$ across the panel of 8 PEL cell lines is shown.

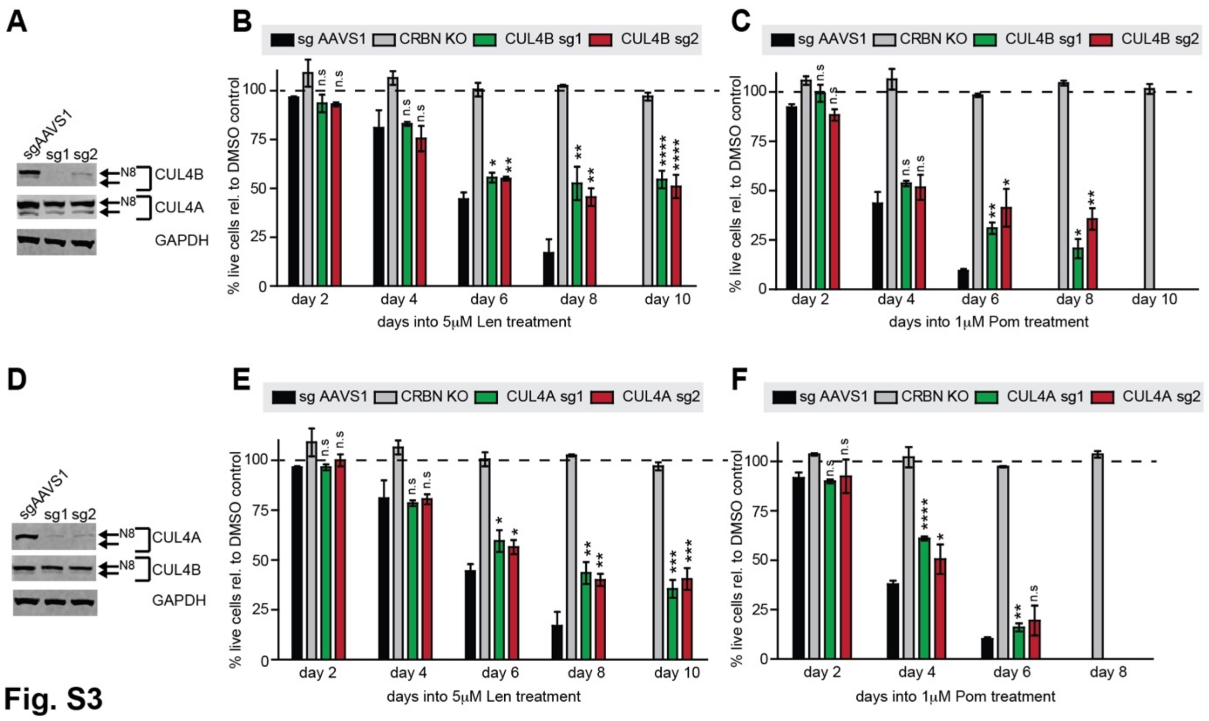


**Figure S3. Inactivation of CUL4A/B confers resistance to LEN.** (A) Representative Western blot analysis confirms efficient CRISPR-induced inactivation of CUL4B in BC-3/Cas9 cells by two independent CUL4B-specific sgRNAs (sg1, sg2). These sgRNAs did not affect CUL4A expression. sgAAVS1 served as negative control guide, GAPDH served as loading control. (B-C) Growth curve analyses of CUL4B-inactivated BC-3/Cas9 cells after treatment with 5µM LEN (B) or 1µM POM (C). BC-3/Cas9 cells transduced with sgAAVS1 were included as a CM-sensitive negative control, while previously described clonal CRBN-inactivated BC-3 cells (CRBN KO) served as a positive control for complete CM resistance. Absolute live cell numbers were normalized to corresponding DMSO vehicle treated cells at each passage (represented by the dotted line). Experiments were run in parallel with those shown in Fig. 3B-C and 4B-C and thus share common negative and positive controls for several or all replicates (n=3, error bars represent SEM). (D) As in A, except that CUL4A-specific sgRNAs were used. (E-F) As in (B-C), except that CUL4A-specific sgRNAs were used. (n=3, error bars represent SEM). Statistical analyses throughout this figure were done by unpaired *t*-tests comparing specified conditions to corresponding sgAAVS1 controls. * $p<0.05$, ** $p<0.01$, *** $p<0.001$, **** $p<0.0001$, n.s. not significant.

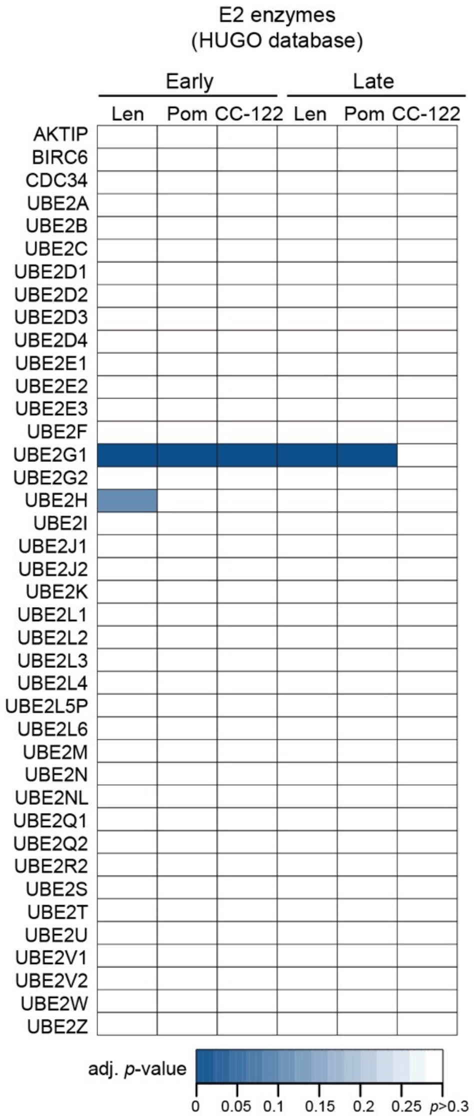


**Figure S4. Gene level sgRNA enrichment of E2 Ubiquitin conjugating enzymes in CM resistance screens.** Heatmap depicting FDR adjusted *p*-value for sgRNA enrichment for all human E2 Ub conjugating enzymes at early or late time points in the CM resistance screens.

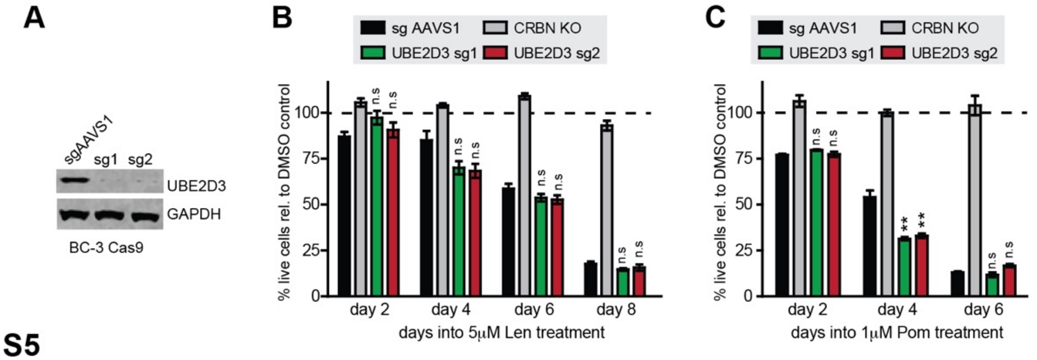


**Fig. S5**

**Figure S5. Inactivation of UBE2D3 in BC-3 cells fails to confer resistance to CMs.** (A) Representative Western blot analysis confirms efficient CRISPR-induced inactivation of UBE2D3 in BC-3/Cas9 cells by two independent UBE2D3-specific sgRNAs (sg1, sg2). sgAAVS1 served as negative control guide, GAPDH served as loading control. (B-C) Growth curve analyses of UBE2D3-inactivated BC-3/Cas9 cells after treatment with 5μM LEN (B) or 1μM POM (C). BC-3/Cas9 cells transduced with sgAAVS1 were included as a CM-sensitive negative control, while previously described clonal CRBN-inactivated BC-3 cells (CRBN KO) served as a positive control for complete CM resistance. Absolute live cell numbers were normalized to corresponding DMSO vehicle treated cells at each passage (represented by the dotted line). Experiments were run in parallel with those shown in Fig. 3B-C and 4B-C and thus share common negative and positive controls for several or all replicates (n=3, error bars represent SEM). Statistical analyses throughout this figure were done by unpaired *t*-tests comparing specified conditions to corresponding sgAAVS1 controls. ** $p<0.01$, n.s. not significant.

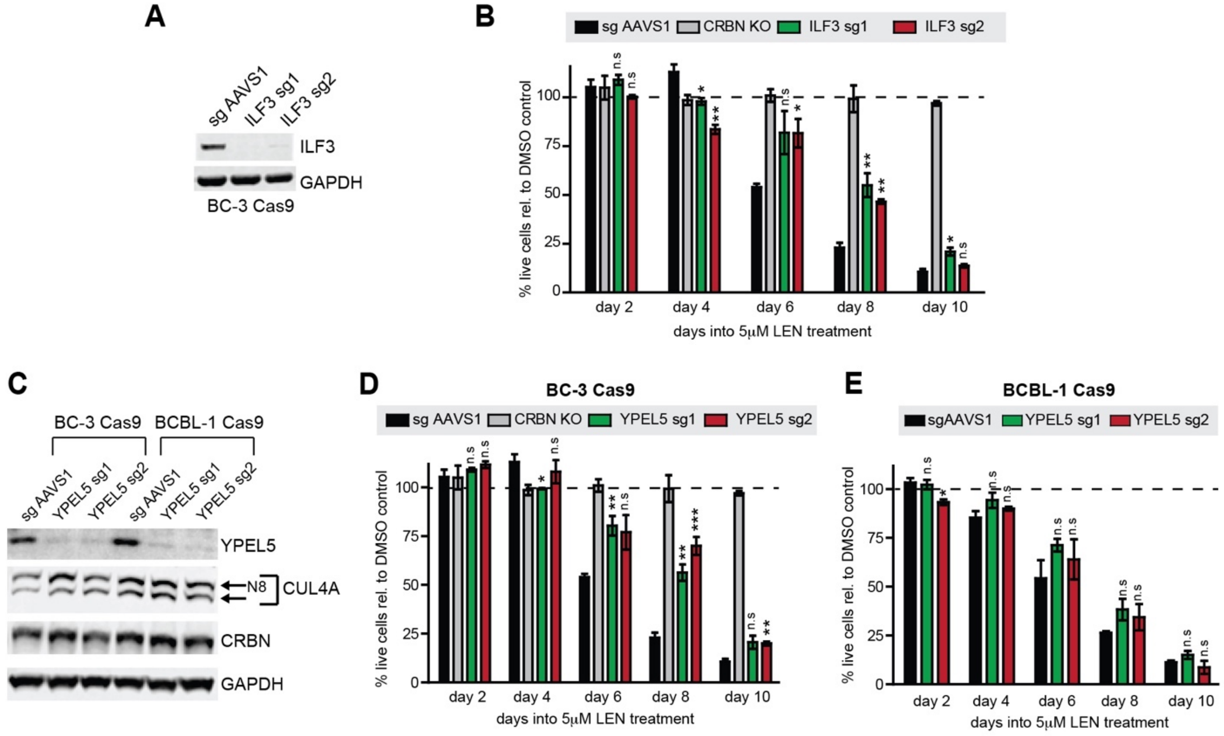


**Figure S6. Inactivation of ILF3 and YPEL5 confers significant resistance to lenalidomide.** (A) Representative Western blot analysis confirms efficient CRISPR-induced inactivation of ILF3 in BC-3/Cas9 cells by two independent ILF3-specific sgRNAs (sg1, sg2). sgAAVS1 served as negative control guide, GAPDH served as loading control. (B) Growth curve analysis of ILF3-inactivated BC-3/Cas9 cells after treatment with 5μM LEN. BC-3/Cas9 cells transduced with sgAAVS1 were included as a CM-sensitive negative control, while previously described clonal CRBN-inactivated BC-3 cells (CRBN KO) served as a positive control for complete CM resistance. Absolute live cell numbers were normalized to corresponding DMSO treated cells at each passage (represented by the dotted line) (n=3, error bars represent SEM). (C) Representative Western blot analysis confirms efficient CRISPR-induced inactivation of YPEL5 in BC-3/Cas9 and BCBL-1/Cas9 cells by two independent YPEL5-specific sgRNAs (sg1, sg2). Membranes were also probed for CUL4A and CRBN, which appeared unaffected. Neddylated (N8) and unneddylated forms of CUL4A are indicated by arrows. sgAAVS1 served as negative control guide, GAPDH served as loading control. (D-E) Growth curve analysis of YPEL5-inactivated BC-3/Cas9 (D) or BCBL-1/Cas9 (E) cells after treatment with 5μM LEN. Cells transduced with sgAAVS1 were included as a CM-sensitive negative control, while previously described clonal CRBN-inactivated BC-3 cells (CRBN KO) served as a positive control for complete CM resistance. Absolute live cell numbers were normalized to corresponding DMSO vehicle treated cells at each passage (represented by the dotted line) (n=3, error bars represent SEM). Statistical analyses throughout this figure were done by unpaired *t*-tests comparing specified conditions to corresponding sgAAVS1 controls. * $p<0.05$, ** $p<0.01$, *** $p<0.001$, n.s. not significant.

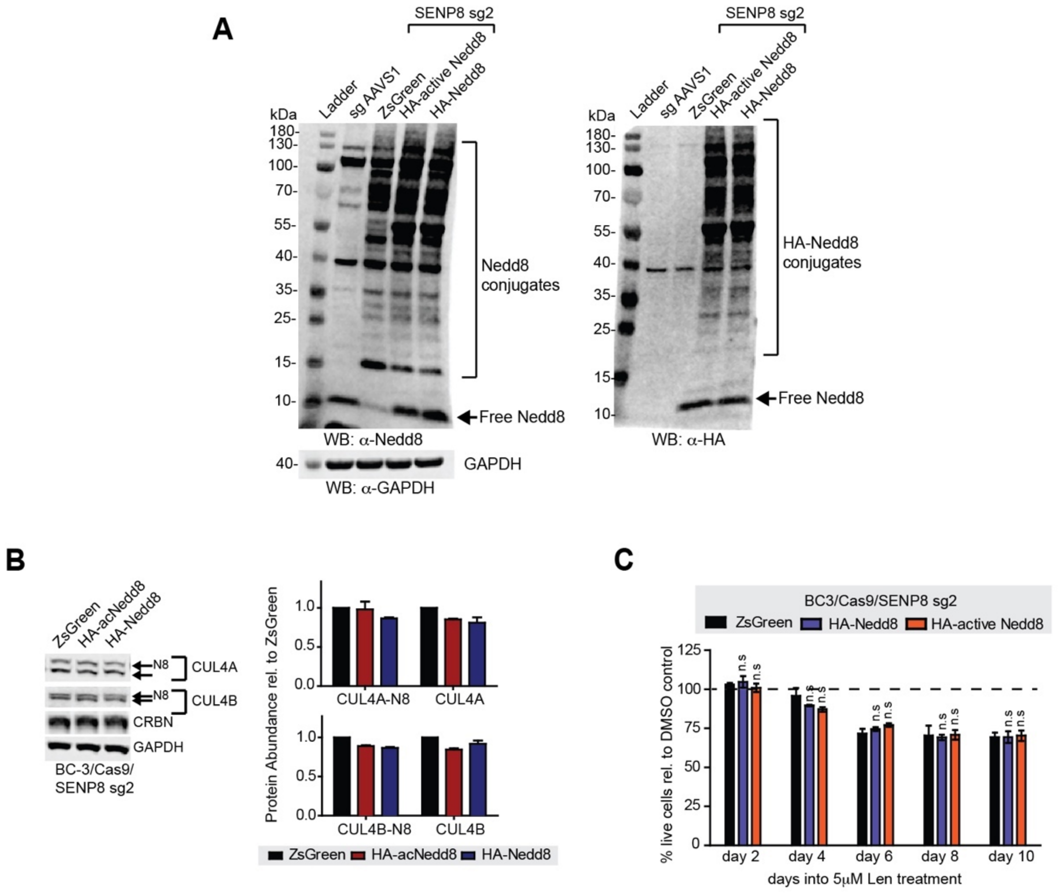


**Figure S7. Nedd8 overexpression fails to restore CUL4A/B neddylation in SENP8 inactivated PEL cells.** (A) Representative Western blot analyses of SENP8-inactivated BC-3/Cas9 cells (using SENP8-specific sg2), which were lentivirally transduced to express HA-tagged precursor or active forms of Nedd8 or ZsGreen. sgAAVS1-transduced cells were included as a control with wild-type neddylation pattern. (B) Lysates described in panel A were subjected to additional Western blot analyses of CUL4A, CUL4B and CRBN expression. Neddylated (N8) and unneddylated forms of CUL4A/B are indicated by arrows. GAPDH was the loading control. The bar graph on the right is a quantitative analysis of neddylated and unneddylated CUL4A/B expression over two replicates. Error bars represent SEM. (C) Growth curve analysis of SENP8-inactivated BC-3/Cas9 cells (using SENP8-specific sg2) overexpressing ZsGreen or HA-tagged precursor or active forms of Nedd8 (see Supplementary Methods) and treated with 5μM LEN or DMSO for the indicated number of days. Live cell counts were expressed relative to those of DMSO-treated cells (n=3, error bars represent SEM). Statistical analyses throughout this figure were done by unpaired *t*-tests comparing specified conditions to corresponding ZsGreen controls. n.s. not significant.

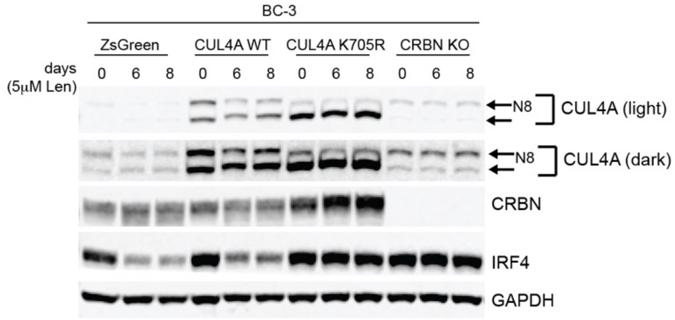


**Figure S8.** Representative Western blot analysis of CUL4A, CRBN and IRF4 at indicated time points confirms a lack of IRF4 downregulation in BC-3 cells expressing CUL4A K705R after treatment with 5μM LEN. Neddylated (N8) and unneddylated forms of CUL4A are indicated by arrows. Samples are matched to growth curve shown in Fig. 6D. GAPDH served as loading control.
